# Structural rearrangement of the intracellular gate of the serotonin transporter induced by Thr276 phosphorylation

**DOI:** 10.1101/2021.10.13.464332

**Authors:** Matthew C. Chan, Erik Procko, Diwakar Shukla

## Abstract

The reuptake of the neurotransmitter serotonin from the synaptic cleft by the serotonin transporter, SERT, is essential for proper neurological signaling. Biochemical studies have shown Thr276 of transmembrane helix 5 is a site of PKG-mediated SERT phosphorylation, which has been proposed to shifts the SERT conformational equlibira to promote inward-facing states, thus enhancing 5HT transport. Recent structural and simulation studies have provided insights into the conformation transitions during substrate transport but have not shed light on SERT regulation via post-translational modifications. Using molecular dynamics simulations and Markov state models, we investigate how Thr276 phosphorylation impacts the SERT mechanism and its role in enhancing transporter stability and function. Our simulations show that Thr276 phosphorylation alters the hydrogen-bonding network involving residues on transmembrane helix 5. This in turn decreases the free energy barriers for SERT to transition to the inward-facing state, thus facilitating 5HT transport. The results provide atomistic insights into *in vivo* SERT regulation and can be extended to other pharmacologically important transporters in the solute carrier superfamily.

## 1 Introduction

The serotonin transporter (SERT, SLC6A4) is responsible for the reuptake of synaptic serotonin (5-hydroxytryptamine, 5HT) from the synapse thereby regulating serotonergic signaling in the brain and elsewhere in the body. SERT, as well as the dopamine transporter (DAT) and norephephine transporter (NET), are members of the sodium-coupled, chloride-dependent monoamine transporters in the neurotransmitter:sodium symporters (NSS) family and the solute carrier (SLC) superfamily (*1*). Members of this family adopt an inverted psudeo-symmetrical architecture consisting of 12 transmembrane (TM) helices commonly known as the LeuT fold(*1*, *2*) (Figure 1A). Transport of neurotransmitters across the neuronal membrane via the NSS family is facilitated by an alternating access mechanism in which these transporters then undergo a series of conformational transitions from an outward-facing (OF) state, where the binding cavity is accessible from the extracellular side, to an occluded (OC) state, and finally an inward-facing (IF) state where the substrates are released into the neuron (Figure 1) (*3*). Reverting back to the outward-facing state involves the efflux of potassium ions in some NSS transporters (*4*).

**Figure 1:**
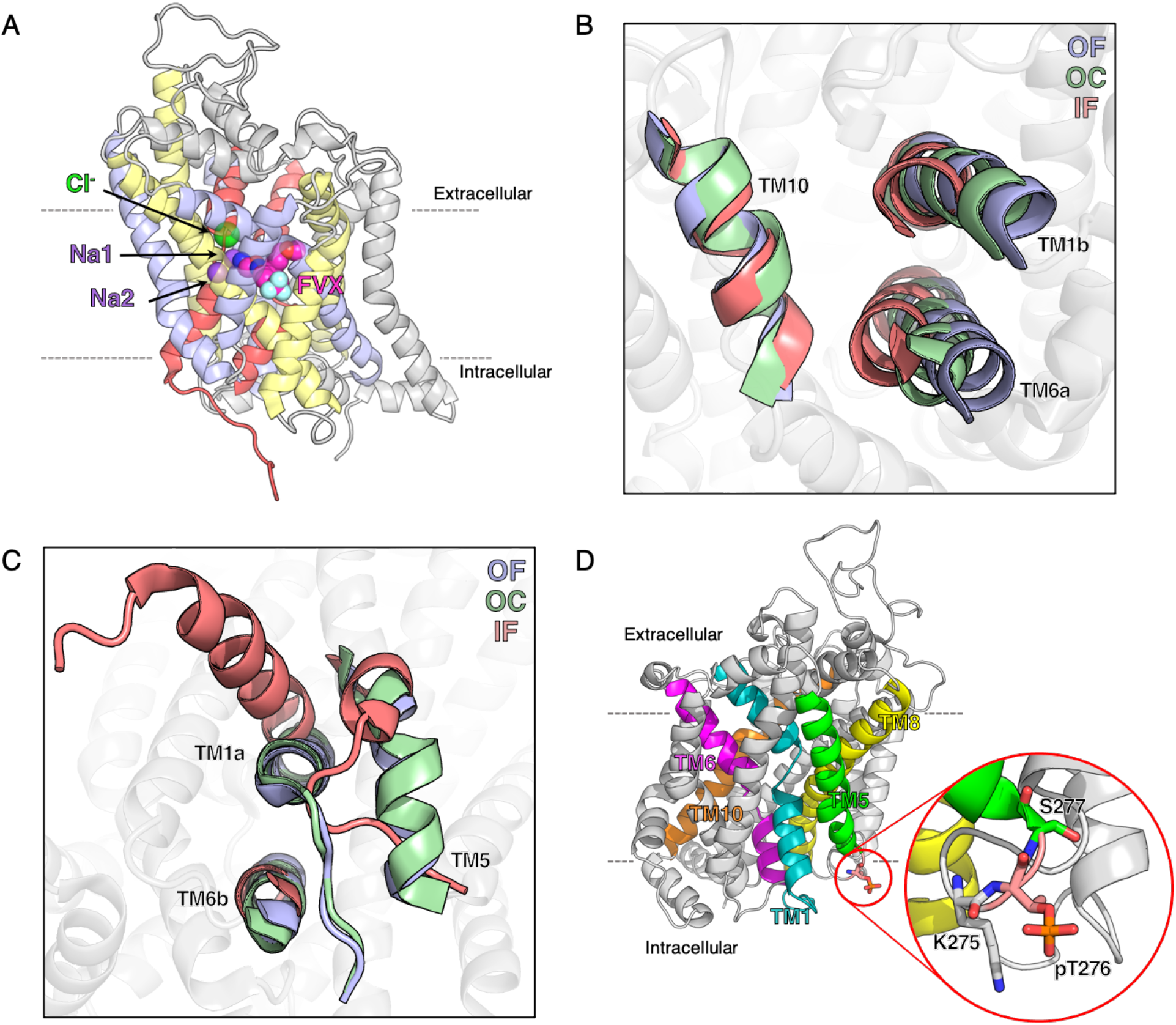
Architecture of the serotonin transporter, SERT. (A) Crystal structure of SERT complexed with inhibitor molecule fluvoxamine resolved in the outward facing conformation (PDB: 6AWP). The sodium and chloride ions resolved in the crystal structure are shown as purple and green spheres, respectively. Fluvoxamine (FVX) bound in the orthosteric site shown in magenta spheres. The two fold architecture of SERT is colored as follows: TM1 and TM6 in red, TM2-5 in light blue, TM7-10 in yellow, TM11-12 in gray. (B, C) SERT viewed from the extracellular plane (B) and intracellular plane (C) showing the conformational transitions of the gating helices involved in the transport process. The cryo-EM structures of SERT resolved in three states (OF (PDB:6DZY): blue, OC (PDB:6DZV): green, IF (PDB:6DZZ): salmon) are overlaid on the SERT-OF structure (gray). (D) The Thr276 phosphorylation site on TM5 is shown as salmon colored sticks. The SERT structure is represented as cartoon with TM 1, 5, 6, 8, and 10 colored in teal, green, magenta, yellow, and orange, respectively.

In the body, SERT is regulated thorough numerous phosphorylation mechanisms that involve protein kinases, phosphatases, receptors, and substrates with implications for transporter expression, stability, trafficking, oligermization, and uptake activity (*5*–*7*). Consequently, improper regulation of transporter function is associated with various physiological complications and psychiatric disorders (*8*–*11*). Increased phosphorylation of SERT by protein kinase C-linked pathways is linked with increased SERT internalization and decreased 5HT-transport activity (*12*, *13*). Upregulation of SERT by protein kinase G (PKG) enhances expression and transport activity (*14*). The psycho-stimulant drug amphetamine increases SERT phosphorylation (*13*) and in DAT, amphetamine-induced phosphorylation of N-terminal residues exhibits a dopamine efflux function (*15*, *16*). Among other transporters in the SLC superfamily, phosphorylation heavily influences transporter function, and thus is a universal mechanism of regulating transporter activity (*17*–*20*).

From a thermodynamics perspective, post-translational modifications (e.g. phosphorylation, glycosylation, lipidation, protonation) may alter the conformational free energy landscape, thus affecting protein stability and/or dynamics (*21*, *22*). The use of molecular dynamics (MD) simulations not only provide an atomistic perspective of complex protein dynamics, but upon sufficient sampling, may allow us to quantify the thermodynamics of functional states and key transition barriers. For example, serine/threonine phosphorylation of protein kinases promotes active-like conformations by stabilizing the dynamics of flexible loops (*23*–*25*). Alternatively, phosphorylation(*24*) and s-glutathionylation(*26*) of the plant kinase BAK1 alter the free energies where inactive states are favored over active-like states. Tyrosine nitration of an abscisic acid plant hormone receptor increases the free energy barriers for ligand binding, thereby preventing receptor activation (*27*). As a final example, glycosylation of SH3 domains promote their folded states due to the presence of bulky side chains that destabilize unfolded states (*28*). Therefore, relating how post-translational modifications affect the protein dynamics and conformational free energy landscape is necessary to understand how protein function is regulated.

In 2007, Ramamoorthy *et al*. identified Thr276 of TM5 to be a site of PKG-mediated SERT phosphorylation (Figure 1D). These observations uncovered essential insights into the *in vivo* regulatory mechanisms of SERT trafficking and transport function via post-translational modification (*29*). It was later characterized by Zhang *et al*., through the binding of conformational selective inhibitors cocaine and ibogaine, that Thr276 phosphorylation directly modulates the conformational equilibra of functional states to enhance 5HT-transport (*30*). Quantum dot studies conducted by Bailey *et al*. further correlated Thr276 phosphorylation with cholesterol depletion in midbrain neurons (*31*). We have previously conducted large-scale MD simulations to characterize the serotonin import process in SERT. We showed how 5HT binding in the orthosteric site reduces the free energy barriers for transition from the outward-facing to inward-facing states, while also stabilizing the inward-facing state to promote substrate import (*32*, *33*). In this current study, we aim to understand the molecular mechanism of Thr276 phosphorylation and its structural consequences on the conformational heterogeneity of SERT. We first preformed MD simulations of SERT bound with inhibitors to provide atomistic details of Zhang *et al*.’s observations (*30*). Next, using our previously collected SERT data as a comparison(*32*), we characterized the dynamics and structural stability of phosphorylated Thr276 SERT using Markov state models. To this extent, we collected over 600 microseconds of MD simulations data using the distributive computing platform Folding@Home (*34*) of phosphorylated Thr276 SERT. Our results show that Thr276 phosphorylation modulates SERT dynamics primarily through the rearrangement of intracellular hydrogen bonding interactions. Consequently, the altered dynamics of the intracellular gating helices reduces the free energy barriers between occluded and inward-facing states and further stabilizes SERT in the inward-facing state for substrate release into the cell.

## 2 Results and discussion

### 2.1 Accessibility of the Thr276 phosphorylation site under inhibitor binding

Structural and computational studies on SERT have revealed that structural transitions from the outward-facing state to the inward-facing state of SERT are initiated by the binding of substrates in the orthosteric pocket which triggers the movement of extracellular gating helices TM1a and TM6b towards the helical scaffold (*32*, *35*, *36*) (Figure 1B). The closure of the extracellular vestibule stabilizes the transporter to allow for solvation of the intra-cellular exit path and the formation of the inward-facing state. The conformation of the inward-facing state is notably associated with the outward motion of TM1a from the helical bundle and unwinding of the cytoplasmic base of TM5 to promote a solvent exposed intracellular vestibule for substrate release (*36*–*39*) (Figure 1C). Multiple studies conducted by the Rudnick group investigated the reactivity of substituted cysteine residues with MTSEA (2-(aminoethyl)methanethiosulfonate hydrobromide) as a measure of SERT accessibility and conformational transitions (*30*, *37*, *40*–*43*). Of these studies, in 2016, Zhang *et al*. used cocaine and ibogaine to influence the conformational equilibra of outward-facing and inward-facing states and investigated the effects of Thr276 phosphorylation on the conformational dynamics and substrate transport mechanism(*30*). They have identified PKG-mediated phosphorylation of Thr276 to occur more readily when SERT is bound with ibogaine as compared to cocaine. As ibogaine stabilizes SERT in an inward-facing state, this allows for TM5 to unwind and promote Thr276 phosphorylation. The cryo-EM structure of SERT bound with ibogaine would later be resolved to depict the unfolded structure of TM5 (*36*).

To provide an atomistic perspective of Zhang *et al*.’s observations, we performed MD simulations of SERT bound with inhibitors at the orthosteric site. Cocaine docked in an outward-facing SERT crystal structure (PDB: 6AWO) or the ibogaine-complexed inward-facing cryo-EM SERT structure (PDB: 6DZZ) were used as the starting structures for simulations (Figure 2B, 2C). The proteins were embedded in a 1-palmitoyl-2-oleoyl-sn-glycero-3-phosphocholine (POPC) lipid bilayer and solvated with 150mM NaCl. Five independent 100 ns long simulations for each SERT-inhibitor complex were performed.

**Figure 2:**
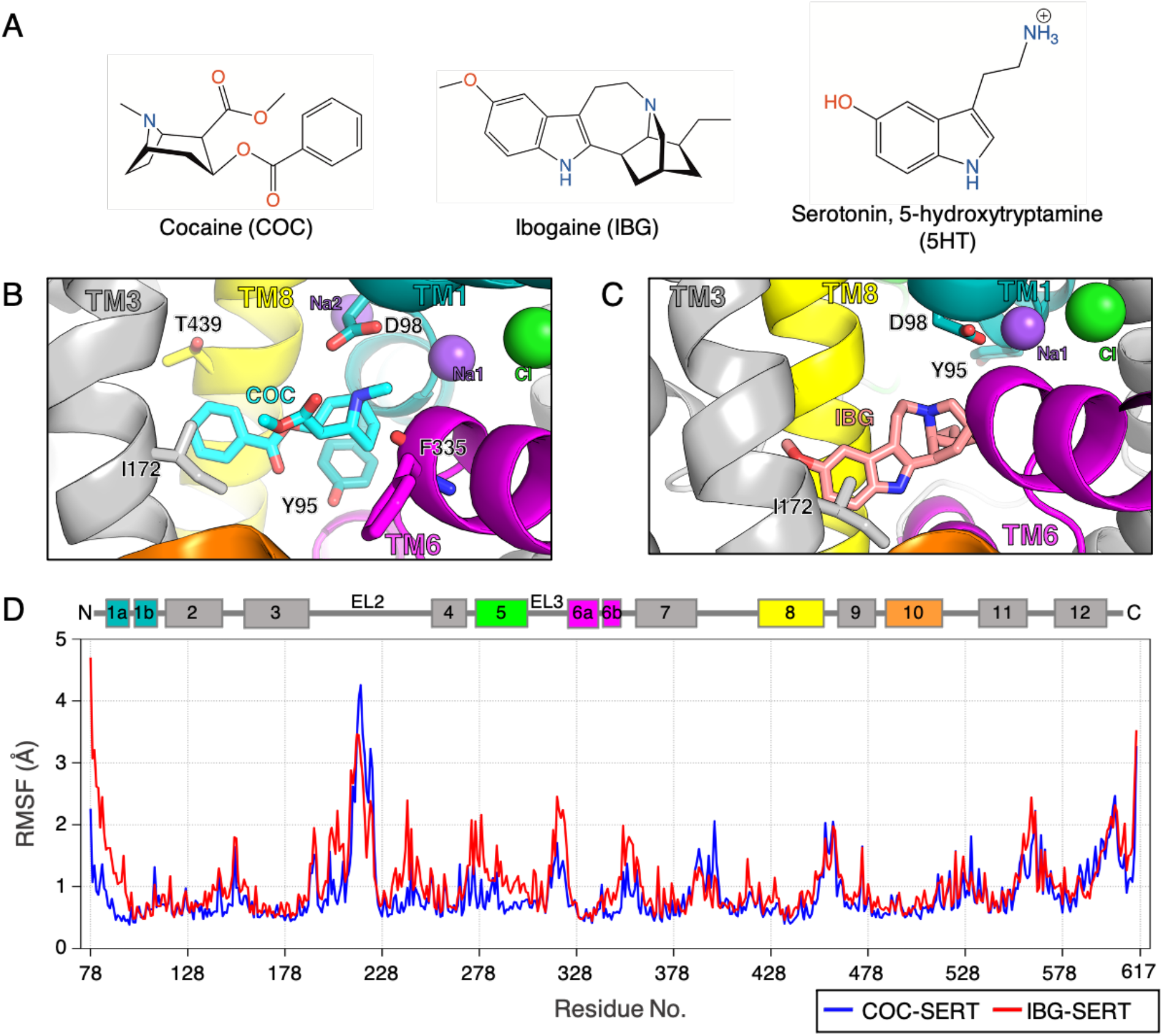
Dynamics of inhibitor-bound SERT. (A) Chemical structures for the conformational selective inhibitors cocaine and ibogaine and the endogenous substrate serotonin (5HT). (B, C) MD snapshots of cocaine (B) and ibogaine (C) bound in the orthosteric site. TM helices 1, 5, 6, 8, and 10 colored in teal, green, magenta, yellow, and orange respectively. (D) Root-mean-square fluctuation (RMSF) of ibogaine-bound SERT (red; in the IF state) and cocaine-bound SERT (blue; in the OF state). The calculated RMSF was averaged over five independent 100 ns simulations.

The simulations show distinct structural characteristics of the respective SERT-inhibitor complex. As expected, the fluctuations of TM1a in the intracellular vestibule are greater when SERT is in the ibogaine-bound inward-facing state versus the cocaine-bound outward-facing state (Figure 2). Additionally, the fluctuations of extracellular loop (EL) 2 are more pronounced in simulations of the SERT-cocaine complex. This observation was also noted in our previous study illustrating the coupled dynamics of EL2 and the opening and closure of the extracellular vestibule (*32*). Most importantly, unwinding of the cytoplasmic base of TM5 in ibogaine-bound SERT promotes greater dynamics of the entire helix and intracellular loop (IL) 2.

Solvent accessible surface area (SASA) measurements show increased solvent exposure of Thr276 in ibogaine-bound SERT simulations as compared to cocaine-bound SERT (Figure 3A). In the inward-facing SERT-ibogaine structure, the outward tilt of TM1a enables Tyr95 to interact with the backbone carbonyl of Thr276 while Tyr350 hydrogen bonds with Gly273. These interactions initially stabilize the unfolded TM5, but after ~20 ns, the hydrogen bonding interactions break and the unfolded TM5 region becomes exposed to the intracellular solvent. Afterwards, Thr276 remains exposed to the solvent, with an average SASA of 82 ± 12 Å^2^ as compared to 26 ± 11 Å^2^ in SERT-cocaine simulations. Furthermore, these observations are consistent with SASA calculations from our previous simulations with the endogenous substrate 5HT (Figure 3B) (*32*). The binding of 5HT enables similar transition to the inward-facing state where unwinding of TM5 further allows Thr276 to be exposed to the cytoplasm. Overall, our observations of the accessibility of the Thr276 phosphorylation site is consistent with the findings presented by Zhang *et al*.

**Figure 3:**
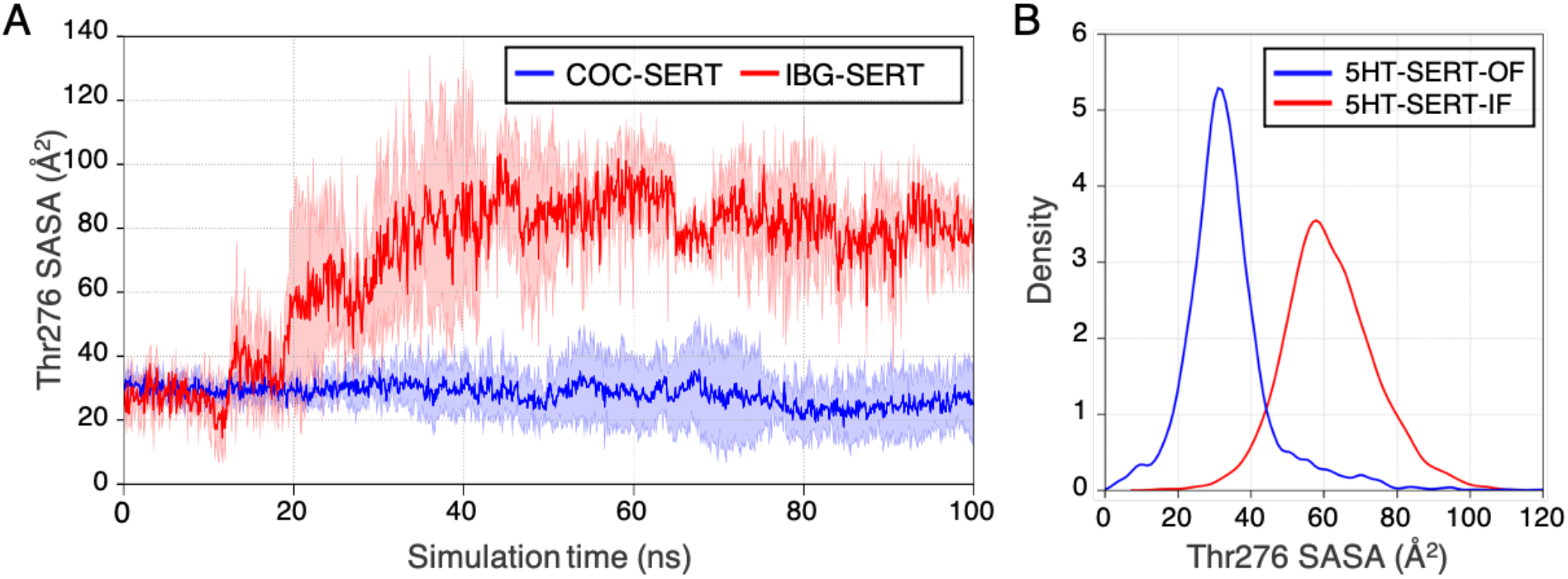
Accessibility of Thr276. (A) Calculated solvent accessible surface area (SASA) of Thr276 for SERT-ibogaine (red) and SERT-cocaine (blue). The calculated SASA was averaged over five independent 100 ns simulations. (B) Density distribution of Thr276 SASA from outward- and inward-facing states from SERT-5HT import simulations (*32*).

### 2.2 Phosphorylation of Thr276 alters the conformational free energy landscape

By projecting the electrostatic potential of the three-dimensional structure of SERT, we observed that phosphorylation of Thr276 affects the intracellular gate and neighboring residues (Figure S1). When closed, there is a positive surface charge at the intracellular gates of de-phosphorylated SERT (dphos-SERT). When Thr276 is phosphorylated, residues surrounding the phosphorylation site become neutralized, thereby potentially altering the dynamics of the intracellular gate during occluded to inward-facing transitions. Given the proximity of Thr276 to the intracellular gating domain, we sought to understand how phosphorylation affects the intrinsic dynamics using MD simulations of phosphorylated Thr276 SERT (pThr276-SERT). To efficiently explore the conformational landscape, we seeded 2,520 independent pThr276-SERT simulations to be conducted on Folding@Home (*34*). The starting structures were selected from a Markov state model (MSM)-weighted distribution of 18 macrostates of the dphos-SERT obtained from our previous study (*32*). An aggregated total of ~630 *μ*s of simulation data were collected and used to construct a MSM (see Methods for details).

Projection of the MSM-weighted simulation data on the axes defined by the extracellular and intracellular gating residues quantifies the relative stability of SERT conformational states (Figure 4). In dphos-SERT simulations, transitions from the occluded state to inward-facing state were rate limiting for substrate import, with free energy barriers of ~2 kcal/mol (Figure 4A) (*32*). Additionally, as compared to outward-facing and occluded states, the formation of inward-facing states in dphos-SERT are relatively less stable. The modification of Thr276 to phosphothreonine exhibits shifts in the free energy barriers of the conformational landscape. Transitions from outward-facing to occluded states retain relatively low free energy barriers. While outward-facing and occluded states remain stable with a relative free energy of ~0-1 kcal/mol, the inward-facing state is further stabilized by ~0.5-1 kcal/mol (Figure 4B). The transitions from occluded to inward-facing are further reduced by ~0.5 kcal/mol as compared to dphos-SERT (Figure 4C), in agreement with the 25% increase in 5HT uptake as experimentally characterized by Zhang *et al*. (*30*).

**Figure 4:**
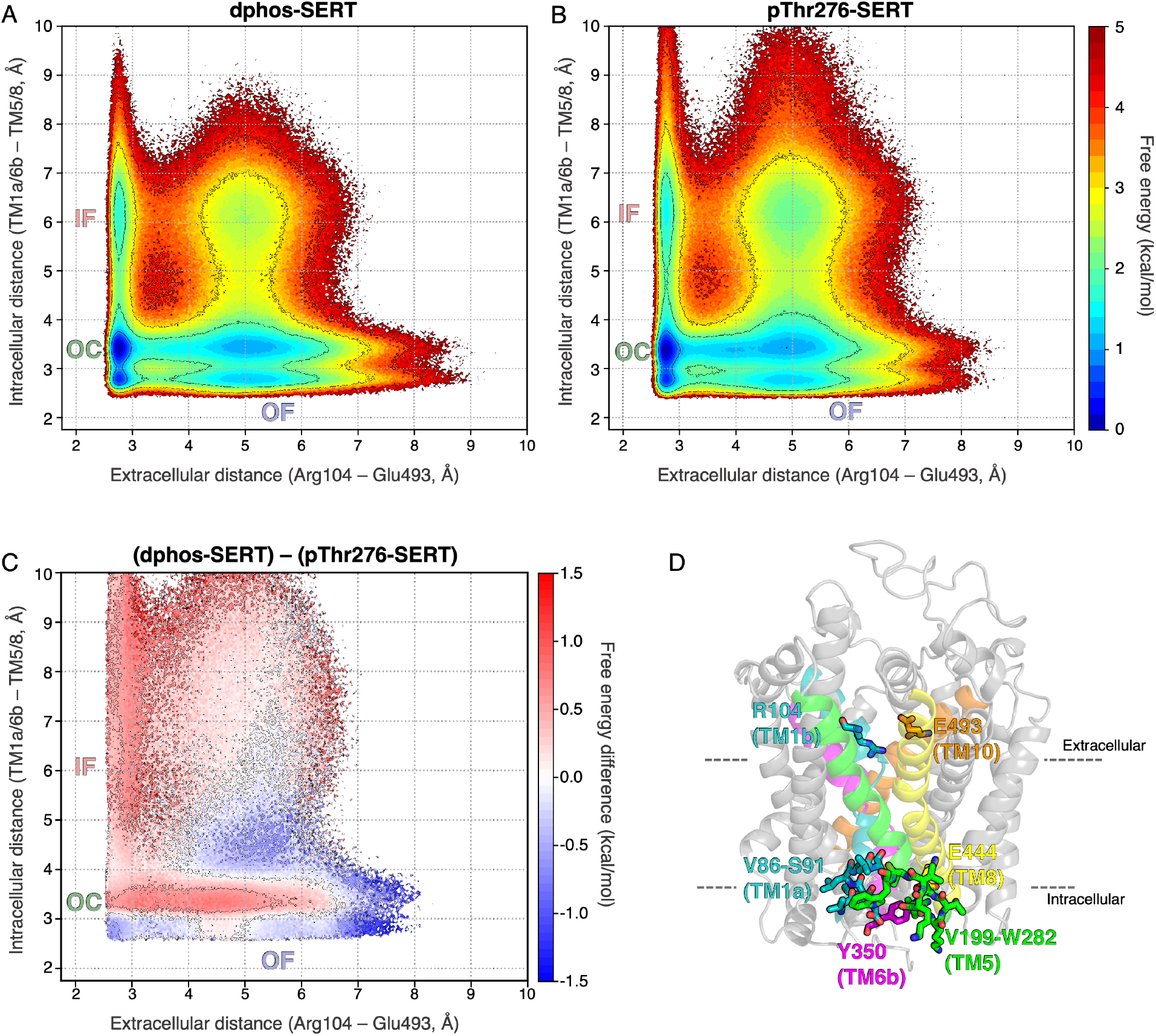
Phosphorylation of Thr276 alters the SERT conformational free energy landscape. (A, B) Conformational free energy landscapes for dphos-SERT (A) and pThr276-SERT (B) projected on the coordinates defined by the extracellular and intracellular gating distances. Simulation data were reweighted by the Markov state model equilibrium probabilities. (C) Difference between the of free energy landscapes of dphos-SERT and pThr276-SERT projected on the same coordinates of the gating distances. (D) Metrics used for the projection of the free energy landscapes. Conformations in red have relatively lower free energy in pThr276-SERT. Extracellular gating distances were defined as the closest heavy atom between Arg104 and Glu493. Intracellular gating distances were calculated between residues of TM1a/TM6b and TM5/TM8.

### 2.3 Rearrangement of the intracellular hydrogen-bonding network

The intracellular gate of SERT is comprised of a number of charged residues on TM1a, TM5, TM6b, and TM8 that form a hydrogen bonding network to stabilize the transporter in outward-facing and occluded states (Figure 5A). These residues are conserved among other monoamine transporters as well as the NSS family. Several studies have highlighted the importance of the intracellular region in the NSS family and its role in the gating mechanism (*32*, *36*, *44*–*46*). The binding of the substrates in the orthosteric site closes the extracellular vestibule thereby initiating the breakage of these electrostatic interactions and promoting transitions to the inward-facing state where an intracellular exit pathway is formed between TM1a and TM5 (Figure 5B).

**Figure 5:**
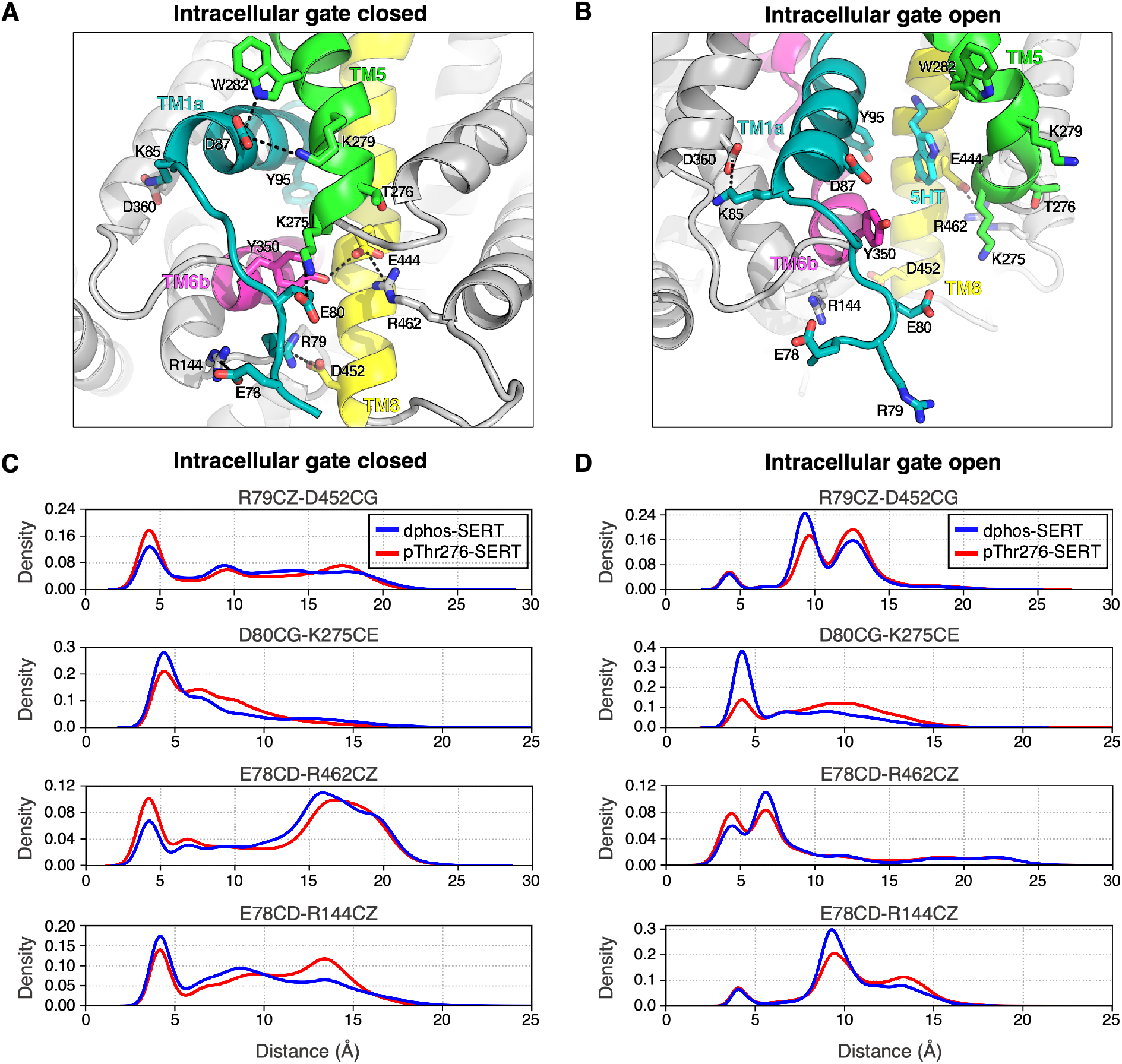
Rearrangement of the intracellular gating residues as a result of Thr276 phosphorylation. (A,B) MD snapshot of dphos-SERT in the occluded (A) and in the inward-facing state (B). Hydrogen bonding pairs that contribute to the closure of the intracellular exit pathway are shown in sticks. (C, D) Distance distribution of select gating residues for 50,000 MD structures for each occluded (C) and inward-facing (D) is shown. Distances calculated from the dphos-SERT MD simulations is represented in blue while pThr276-SERT in red. See Figure S2 for more distance distributions of intracellular residue pairs.

MD simulations of pThr276-SERT shows that most of the intracellular interactions are formed but with slight deviations of the distance distributions as compared to the dphos-SERT simulations (Figure 5C, 5D). We observed that the phosphorylation of Thr276 disrupts the hydrogen bonding interactions of residues on TM5, most notably Lys275, Lys279, and Trp282. The interactions are critical in stabilizing TM5 with TM1a while the intracellular gate is closed. When comparing the occluded structures from simulations, the distances for pThr276-SERT intracellular residue pairs exhibit a broader distribution suggesting overall weaker interactions. For the Asp80-Lys275 pair, the distance between these residues increases in pThr276-SERT simulations, especially when in inward-facing states (Figure 5D, S2). The electrostatic interactions Arg79-Asp452, Glu78-Arg462, and Glu78-Lys275 are more prevalent in pThr276-SERT as compared to dphos-SERT (Figure S2). Given the increased flexibility of the N-terminal tail, residues on the N-terminus may compensate for weaker interactions of TM1a and TM5 when Thr276 is phosphorylated and retain stable outward-facing and occluded states for substrate binding.

The helical structure of TM5 is regulated by the hydrogen bonding interactions between the side chains of Thr276 and Ser277 with the backbone carbonyl of Ser269 on TM4 (Figure 6A). Upon conformational transitions to the inward-facing state, these interactions are severed resulting in the unwinding of the cytoplasmic base of TM5. The addition of the phosphate to Thr276 not only presents a negative surface charge but also prevents Thr276 from being the hydrogen bond donor. Projection of the simulation data on the coordinates defined by the distance of the Thr276 side chain with the backbone carbonyl of Ser269 verses the average helical content of TM5 shows that this interaction is not maintained in pThr276-SERT and allows for greater unfolding of TM5 (Figure 6B). The side chain of Ser277 makes alternate interactions with Ser269 and Glu444 in dphos-SERT simulations. However, in pThr276-SERT, we observed Ser277 maintains its interaction with Ser269, but not with Glu444. This hydrogen bond rearrangement in pThr276-SERT is a result of the outward bend of the pThr276 residue due to its larger and negatively charged sidechain. As a result, TM5 is unable to maintain its helical structure. Furthermore, the structural consequence of Thr276 phosphorylation results in nearby charged residues to heavily interact with the phosphate group of pThr276, thereby weakening the interactions of TM5 (Figure 6C). Due to the flexible nature of the N-terminus, Arg79, which typically interacts with Asp452, forms interactions with the Thr276 phosphate group. Furthermore, the adjacent residue Lys275 may also interact with the phosphate group. These additional interactions may destabilize the intracellular gates and decrease the free energy barriers for transition to the inward-facing state.

**Figure 6:**
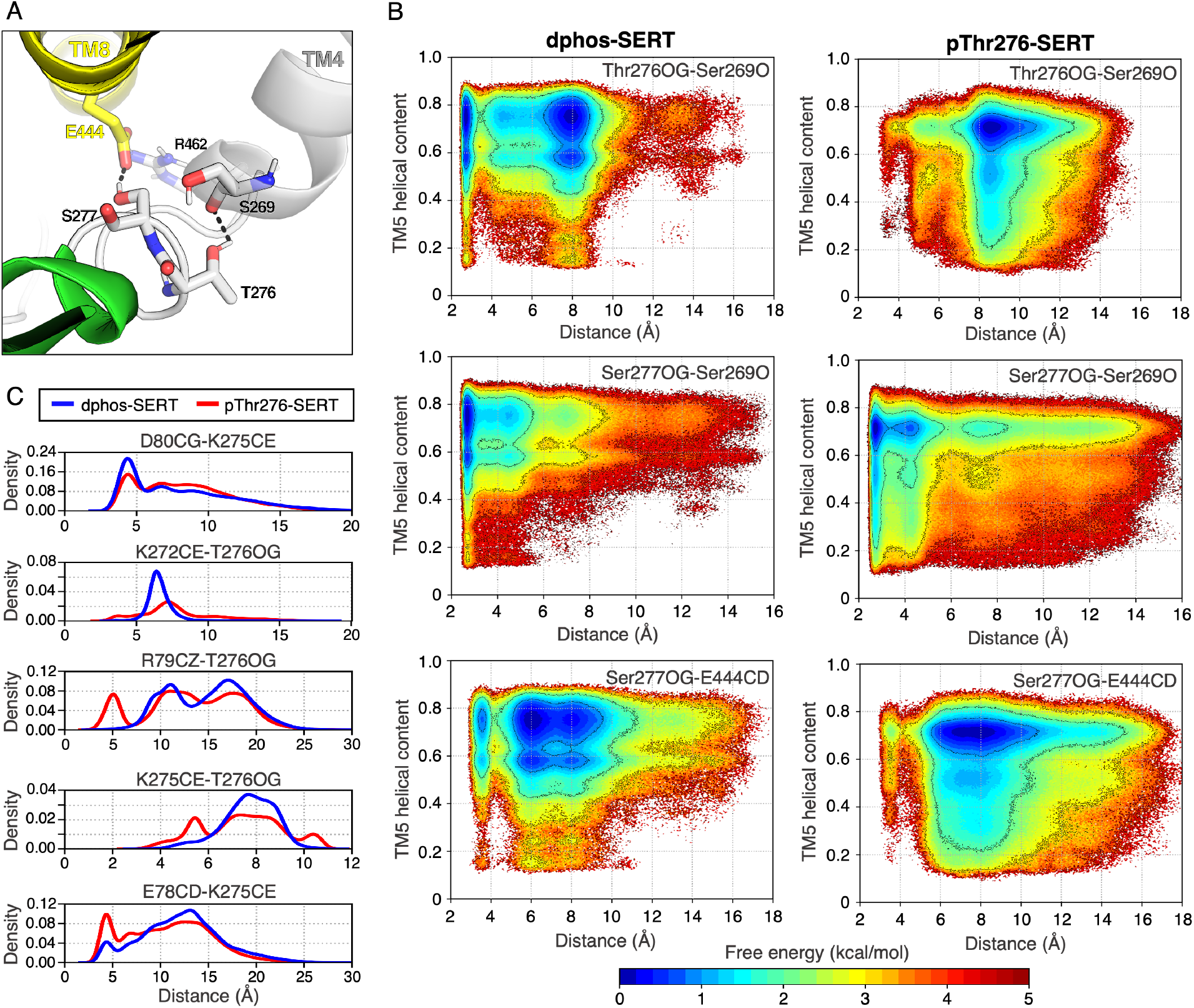
Thr276 phosphorylation further stabilizes the unwinding of TM5 during conformational transitions. (A) MD snapshot of the hydrogen bond arrangement to maintain the helical fold of TM5. The side chain of Thr276 interacts with the backbone carbonyl of Ser269 (TM4), while Ser277 hydrogen bonds with Glu444. (B) Free energy landscapes comparing the helical content of TM5 and hydrogen bonds identified in panel A. Helical content was measured for residues 273 to 280. Phosphorylation of Thr276 not only disrupts the hydrogen bonds formed by Ser269, but also further increases the unfolding of cytoplasmic base of TM5. (C) Shifts in the distance distribution of hydrogen bonds involving residues neighboring Thr276. Distances calculated from the dphos-SERT MSM are represented in blue while pThr276-SERT distance are in red.

Understanding the molecular regulation of neurotransmitter transporters is vital for studying normal neurological function in the brain and developing therapeutics to treat various psychiatric disorders. The observations presented in this study provide an atomistic perspective into the mechanism of regulating SERT conformational dynamics by Thr276 phosphorylation. Using adaptive sampling and Markov state models to explore the SERT conformational space, we find that phosphorylated Thr276 results in the rearrangement of the intracellular hydrogen bonding network, particularly residues involving TM5. These altered interactions consequently decrease the free energy barriers for occluded to inward-facing transitions. Furthermore, inward-facing states are further stabilized to allow for substrate release. The results obtained in this work alongside previously conducted experimental studies of Thr276 phosphorylation demonstrate how a phosphorylation event regulates SERT function through altering the conformational equilibria of outward-facing and inward-facing states. Naturally occurring coding variants in human SERT have been shown to alter transporter regulation through changes in PKG/p38 mitogen-activated protein kinase-linked pathways (*47*). In particular, SERT containing the allelic variant Ala56 (normally Gly56 in wild-type) is subjected to hyperphosphorylation under basal conditions (*48*) and suggested to bias SERT in an outward-facing state (*49*). Additionally, the allelic variant Asn605 (normally Lys605 in wild-type) has been proposed to influence SERT in a similar manner as Ala56 (*49*). Other identified phosphorylation sites in SERT may affect overall transporter stability, including but not limited to protein trafficking, expression, and substrate uptake (*50*, *51*).

Aside from phosphorylation, other regulatory mechanisms of the NSS family have been extensively studied through computational and experimental techniques and provide a synergistic approach to characterize *in vivo* transporter regulation. Sterol molecules such as cholesterol have been shown to participate in an inhibitory mechanism among these transporters. A cholesterol molecule wedged between TM1a, TM5, and TM7 was resolved in the crystal structure of the dopamine transporter(*52*). Further investigation through course grain MD simulations shows that this specific cholesterol site inhibits the outward motion of TM1a, thereby locking the transporter in outward-facing states (*53*). Biochemical and computational studies also support a similar mechanism of cholesterol inhibition in SERT (*54*). Phosphatidylinositol 4,5-biphosphate (PIP2)-mediated interactions with the N-terminal residues of DAT have been shown to influence transport function (*55*). Molecular modeling of DAT further shows the electrostatic interactions of PIP2 to promote the opening of the intracellular exit vestibule (*56*). Oligomerization of NSS transporters has been implicated in membrane trafficking and transporter regulation (*57*–*60*). Despite various biochemical and computational studies investigating the effects of oligomerization (*61*–*63*), there is no clear consensus of the oligomeric interface in the NSS family. Furthermore, how the transport function is affected by oligomerization remains unclear. As this work focused on the conformational transitions associated with the substrate import process, how phosphorylated Thr276 affects SERT reverting back from inward-facing to outward-facing transition states remains unknown. A potential potassium binding site remains ambiguous, but it has been shown through biophysical experiments that potassium favors an inward-facing-like state (*64*, *65*). Further studies may investigate how the altered interactions formed due to phosphorylated Thr276 affect the closure of the inward-facing state and transitions to outward-facing in the presence of potassium.

## 3 Methods

### 3.1 Inhibitor bound SERT simulation setup

For the cocaine bound simulations, an outward facing SERT crystal structure with the sertraline bound at the orthosteric site (PDB: 6AWO) (*66*) was used as the starting structure for docking simulations. Thermostable mutations Ala218, Ser439, Ala554, and Ala580 were reverted to the wild type residues, Ile218, Thr439, Cys554, and Cys580, respectively. The two Na^+^ ions and single Cl^-^ ion that were resolved in the crystal structure were retained. The sertraline and cholesterol molecules were removed. Cocaine was then docked into the orthosteric cavity using AutoDock Vina 1.1.2 (*67*). PDBQT files for the outward-facing-SERT and protonated cocaine molecules were constructed using the AutoDock python utility scripts. The grid center of the orthosteric binding site was chosen based on the structural alignment of the *Drosophila* dopamine transporter complexed with cocaine (PDB: 4XP4) (*68*). The grid search space was chosen as a 10 Å x 10 Å x 10 Å box centered at the grid center. The default united-atom scoring function implemented in AutoDock Vina was used to obtain docked ligand configurations. When aligned with 4XP4, the RMSD of the docked cocaine molecule was 0.991 Å. The cocaine docked SERT model was then embedded in a homogeneous 1-palmitoyl-2-oleoyl-sn-glycero-3-phosphocholine (POPC) lipid bilayer, solvated with TIP3P water molecules (*69*). 150mM NaCl was added to neutralize the system. Terminal chains were capped with acetyl and methyl amide groups. Glu508 was modeled as the protonated form. A disulfide bridge was modeled between Cys200 and Cys209. Amber ff14SB force field (*70*) was used to parameterize the system. Force field parameters for cocaine were derived using the antechamber (*71*) module of Amber (*72*).

For ibogaine bound simulations, the inward-facing SERT cryo-EM structure complexed with ibogaine (PDB: 6DZZ) (*36*) was used as the starting structure. The protein was embedded in a POPC lipid bilayer and solvated in TIP3P water molecules (*69*) and 150mM NaCl using CHARMM-GUI (*73*). Terminal residues were capped with acetyl and methyl amide groups. Glu508 was modeled as the protonated form. A disulfide bridge was modelled between Cys200 and Cys209. A Cl^-^ and Na^+^ ion were fitted to the Cl^-^ and Na1 site, respectively, based on SERT crystal structures(*36*). As the ibogaine parameters were derived using the CHARMM force field, we parameterized the remainder of the system using the CHARMM36m force fields (*74*). The CHARMM topology files were then converted to Amber format using the chamber module of the parmed program (https://github.com/ParmEd/ParmEd).

### 3.2 Inhibitor bound SERT simulation details

Both cocaine and ibogaine bound SERT simulations were performed using the Amber18 MD package under constant NPT conditions, periodic boundary conditions, and integration timestep of 2 femtosecond. System temperature (300K) was maintained with Langevian dynamics and a 1 picosecond^−1^ damping coefficient. Pressure (1 atm) was maintained with the Monte Carlo barostat with an update interval every 100 steps. Bonds involving hydrogen atoms were constrained using the SHAKE algorithm (*75*). Electrostatics were treated with the particle mesh Ewald method(*76*) and a 10 Å distance cutoff was used to treat nonbonded interactions. Each system was first minimized for 20,000 steps using the conjugate gradient method and then heated to 300K while the protein was constrained. Afterwards, the unrestrainted systems were equilibrated for 50 ns prior to production runs. Five independent simulation runs of 100 ns were performed using the GPU accelerated *pmemd* module of Amber18 (*77*).

### 3.3 Phosphorylated Thr276-SERT simulation on Folding@Home

Our previous study investigated the dynamics of 5HT import of wild-type SERT (*32*). These simulations consisted of 1 SERT protomer (residues 76-616) embedded in a POPC lipid bilayer, solvated with 150mM NaCl and 1 5HT molecules in TIP3P water (*69*). Terminal chains were capped with acetyl and methyl amide groups. Glu508 was modeled as the protonated form. A disulfide bridge was modeled between Cys200 and Cys209. The simulation data previously obtained was used to construct a Markov state model (MSM).

The starting structures for pThr276-SERT simulations were obtained by randomly selecting 70 structures from 18 macrostates of wild-type SERT MSM. For each structure, 2 replicates with different initial random velocities were created, totaling 2,520 independent MD simulations. Thr276 was modified to phosphothreonine using tleap. Additional Na^+^ ions were added to the simulation box to neutralize the system. Each system was prepared using OpenMM 7.4.1 (*78*) and parameterized with an OpenMM ForceField using the Amber ff14SB (*70*) and GAFF force field (*79*). Simulations were performed under periodic boundary conditions and NPT ensemble. The mass of hydrogen atoms and connected atoms were repartitioned according to Hopkins *et al*. (*80*). The Langevin integrator using a timestep of 4 fs, temperature of 300K, and collision rate of 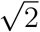 ps^−1^ was used for Langevin dynamics. Pressure (1 atm) was maintained using the Monte Carlo Membrane Barostat with an update frequency of 100 steps. Nonbonded forces were calculated using the particle mesh Ewald method (*76*) with a 10 Å distance cutoff. The resulting OpenMM system and integrator file were serialized to XML format for Folding@Home.

Production simulations for the 2,520 pThr276-SERT systems were conducted on Folding@Home using a simulation core based on OpenMM 7.4.2 (*34*, *78*). A maximum of 250 ns was collected for each system, totaling ~630*μ*s of aggregated simulation data. Simulation snapshots were saved every 100 ps during production runs using mixed precision.

### 3.4 Markov State Models

MSMs are a statistical approach in which the simulation data are discretized into kinetically relevant states and a transition probabilities between each state are calculated. The resulting outcome of the MSM is a kinetic model in which long timescale protein dynamics can be characterized (*81*, *82*). MSMs have been extensively employed to study protein folding, ligand binding and conformational change processes(*83*–*86*). However, there are only few examples of the application of MSMs to membrane transporter proteins(*32*, *33*, *87*–*91*). Here, we employ MSMs to compare the conformational ensemble of the phosphorylated and unphosophorylated SERT to obtain the thermodynamic and kinetic differences responsible for the shift in the conformational equilibrium upon phosphorylation. Finally, theoretical frameworks such as transition path theory (TPT) are used along with MSMs to identify the highest flux pathways and bottlenecks associated with the substrate transport process(*92*).

Trajectories were processed using the CPPTRAJ module of AmberTools (*93*) and MD-Traj Python library (*94*). All pThr276-SERT simulation data were used to construct a Markov state model (MSM) using the pyEMMA Python library (*95*). To maintain consistency among the wild-type SERT and phosphorylated SERT MSM, we used the same 16 residue-residue pair distances along the permeation pathway and z-components of the substrates as described in our previous study. The number of clusters and time-independent components (tICs) were optimized by maximizing the VAMP1 score, or sum of the eigen-values of the transition matrix. The phosphorylated SERT MSM was constructed using 500 clusters, 2 tICs, and a Markovian lag time of 12 ns (Figure S3). Structures extracted from MSM clusters were visualized using Visual Molecular Dynamics (VMD) (*96*) and PyMOL (Schrödinger, LLC). The standard error of the free energy landscapes was calculated by bootstrapping with constructing the MSM with 80% of the trajectory set randomly selected for 500 independent samples (Figure S4). The constructed MSM was further validated using the Chapman-Kolmogorov test performed on 5 macrostates (Figure S5).

## Supporting information

Supplementary Methods, Images, Tables and Results

## Acknowledgement

This work is supported by NSF Early Career Award by NSF MCB 18-45606 to DS and R21 MH113155 from NIMH to EP. This research is part of the Blue Waters sustained-petascale computing project, which is supported by the National Science Foundation (awards OCI-0725070 and ACI-1238993) the State of Illinois, and as of December, 2019, the National Geospatial-Intelligence Agency. Blue Waters is a joint effort of the University of Illinois at Urbana-Champaign and its National Center for Supercomputing Applications. The authors thank Folding@Home donors for computational resources. Authors thank Zhiyu Zhao, Po-Chao Wen, and Emad Tajkhorshid from University of Illinois at Urbana-Champaign for providing parameters for protonated ibogaine.

## References

1. Rudnick, G., Krämer, R., Blakely, R. D., Murphy, D. L., and Verrey, F. (2014) The SLC6 transporters: perspectives on structure, functions, regulation, and models for transporter dysfunction. Pflügers Archiv-European Journal of Physiology 466, 25–42.

2. Forrest, L. R. (2015) Structural symmetry in membrane proteins. Annual Review of Biophysics 44, 311–337.

3. Mitchell, P. (1957) A general theory of membrane transport from studies of bacteria. Nature 180, 134–136.

4. Nelson, P. J., and Rudnick, G. (1979) Coupling between platelet 5-hydroxytryptamine and potassium transport. J Biol Chem 254, 10084–10089.

5. Baudry, A., Pietri, M., Launay, J.-M., Kellermann, O., and Schneider, B. (2019) Multi-faceted Regulations of the Serotonin Transporter: Impact on Antidepressant Response. Frontiers in Neuroscience 13.

6. Ramamoorthy, S., Shippenberg, T. S., and Jayanthi, L. D. (2011) Regulation of monoamine transporters: Role of transporter phosphorylation. Pharmacology & therapeutics 129, 220–238.

7. Cooper, A., Woulfe, D., and Kilic, F. (2019) Post-translational modifications of serotonin transporter. Pharmacological Research 140, 7–13.

8. Ozaki, N., Goldman, D., Kaye, W., Plotnicov, K., Greenberg, B., Lappalainen, J., Rudnick, G., and Murphy, D. (2003) Serotonin transporter missense mutation associated with a complex neuropsychiatric phenotype. Molecular psychiatry 8, 933–936.

9. Hansen, F. H. et al. (2014) Missense dopamine transporter mutations associate with adult parkinsonism and ADHD. Journal of Clinical Investigation 124, 3107–3120.

10. Kitzenmaier, A., Schaefer, N., Kasaragod, V. B., Polster, T., Hantschmann, R., Schindelin, H., and Villmann, C. (2019) The P429L loss of function mutation of the human glycine transporter 2 associated with hyperekplexia. European Journal of Neuroscience 50, 3906–3920.

11. Prasad, H. C., Zhu, C.-B., McCauley, J. L., Samuvel, D. J., Ramamoorthy, S., Shelton, R. C., Hewlett, W. A., Sutcliffe, J. S., and Blakely, R. D. (2005) Human serotonin transporter variants display altered sensitivity to protein kinase G and p38 mitogen-activated protein kinase. Proceedings of the National Academy of Sciences 102, 11545–11550.

12. Ramamoorthy, S., Giovanetti, E., Qian, Y., and Blakely, R. D. (1998) Phosphorylation and regulation of antidepressant-sensitive serotonin transporters. Journal of Biological Chemistry 273, 2458–2466.

13. Ramamoorthy, S., and Blakely, R. D. (1999) Phosphorylation and sequestration of serotonin transporters differentially modulated by psychostimulants. Science 285, 763–766.

14. Steiner, J. A., Carneiro, A. M. D., Wright, J., Matthies, H. J., Prasad, H. C., Nicki, C. K., Dostmann, W. R., Buchanan, C. C., Corbin, J. D., Francis, S. H., and Blakely, R. D. (2009) cGMP-dependent protein kinase Ialpha associates with the antidepressant-sensitive serotonin transporter and dictates rapid modulation of serotonin uptake. Molecular Brain 2, 26.

15. Khoshbouei, H., Sen, N., Guptaroy, B., Johnson, L., Lund, D., Gnegy, M. E., Galli, A., and Javitch, J. A. (2004) N-terminal phosphorylation of the dopamine transporter is required for amphetamine-induced efflux. PLoS Biol 2, e78.

16. Foster, J. D., Yang, J.-W., Moritz, A. E., ChallaSivaKanaka, S., Smith, M. A., Holy, M., Wilebski, K., Sitte, H. H., and Vaughan, R. A. (2012) Dopamine transporter phosphorylation site threonine 53 regulates substrate reuptake and amphetamine-stimulated efflux. Journal of Biological Chemistry 287, 29702–29712.

17. Seo, Y. A., Kumara, R., Wetli, H., and Wessling-Resnick, M. (2016) Regulation of divalent metal transporter-1 by serine phosphorylation. Biochemical Journal 473, 4243–4254.

18. Minematsu, T., and Giacomini, K. M. (2011) Interactions of Tyrosine Kinase Inhibitors with Organic Cation Transporters and Multidrug and Toxic Compound Extrusion Proteins. Molecular Cancer Therapeutics 10, 531–539.

19. Annaba, F., Sarwar, Z., Gill, R. K., Ghosh, A., Saksena, S., Borthakur, A., Hecht, G. A., Dudeja, P. K., and Alrefai, W. A. (2012) Enteropathogenic Escherichia coli inhibits ileal sodium-dependent bile acid transporter ASBT. American Journal of Physiology-Gastrointestinal and Liver Physiology 302, G1216–G1222.

20. Sprowl, J. A. et al. (2016) A phosphotyrosine switch regulates organic cation transporters. Nature Communications 7.

21. Millar, A. H., Heazlewood, J. L., Giglione, C., Holdsworth, M. J., Bachmair, A., and Schulze, W. X. (2019) The Scope, Functions, and Dynamics of Posttranslational Protein Modifications. Annual Review of Plant Biology 70, 119–151.

22. Schönichen, A., Webb, B. A., Jacobson, M. P., and Barber, D. L. (2013) Considering Protonation as a Posttranslational Modification Regulating Protein Structure and Function. Annual Review of Biophysics 42, 289–314.

23. Kuzmanic, A., Sutto, L., Saladino, G., Nebreda, A. R., Gervasio, F. L., and Orozco, M. (2017) Changes in the free-energy landscape of p38*α* MAP kinase through its canonical activation and binding events as studied by enhanced molecular dynamics simulations. eLife 6.

24. Moffett, A. S., and Shukla, D. (2020) Structural Consequences of Multisite Phosphorylation in the BAK1 Kinase Domain. Biophysical Journal 118, 698–707.

25. Jonniya, N. A., Sk, M. F., and Kar, P. (2019) Investigating Phosphorylation-Induced Conformational Changes in WNK1 Kinase by Molecular Dynamics Simulations. ACS Omega 4, 17404–17416.

26. Moffett, A. S., Bender, K. W., Huber, S. C., and Shukla, D. (2017) Allosteric Control of a Plant Receptor Kinase through S-Glutathionylation. Biophysical Journal 113, 2354–2363.

27. Shukla, S., Zhao, C., and Shukla, D. (2019) Dewetting controls plant hormone perception and initiation of drought resistance signaling. Structure 27, 692–702.

28. Shental-Bechor, D., and Levy, Y. (2008) Effect of glycosylation on protein folding: a close look at thermodynamic stabilization. Proceedings of the National Academy of Sciences 105, 8256–8261.

29. Ramamoorthy, S., Samuvel, D. J., Buck, E. R., Rudnick, G., and Jayanthi, L. D. (2007) Phosphorylation of threonine residue 276 is required for acute regulation of serotonin transporter by cyclic GMP. Journal of Biological Chemistry 282, 11639–11647.

30. Zhang, Y.-W., Turk, B. E., and Rudnick, G. (2016) Control of serotonin transporter phosphorylation by conformational state. Proceedings of the National Academy of Sciences 113, E2776–E2783.

31. Bailey, D. M., Catron, M. A., Kovtun, O., Macdonald, R. L., Zhang, Q., and Rosenthal, S. J. (2018) Single quantum dot tracking reveals serotonin transporter diffusion dynamics are correlated with cholesterol-sensitive threonine 276 phosphorylation status in primary midbrain neurons. ACS chemical neuroscience 9, 2534–2541.

32. Chan, M., Selvam, B., Young, H., Procko, E., and Shukla, D. (2020) The Substrate Import Mechanism of the Human Serotonin Transporter. bioRxiv doi: 10.26434/chem-rxiv.9922301.

33. Young, H. J., Chan, M., Selvam, B., Szymanski, S. K., Shukla, D., and Procko, E. (2021) Deep Mutagenesis of a Transporter for Uptake of a Non-Native Substrate Identifies Conformationally Dynamic Regions. bioRxiv doi: 10.1101/2021.04.19.440442.

34. Shirts, M., and Pande, V. S. (2000) Screen savers of the world unite! Science 290, 1903–1904.

35. Koldsø, H., Noer, P., Grouleff, J., Autzen, H. E., Sinning, S., and Schiøtt, B. (2011) Unbiased Simulations Reveal the Inward-Facing Conformation of the Human Serotonin Transporter and Na+ Ion Release. PLoS Computational Biology 7, e1002246.

36. Coleman, J. A., Yang, D., Zhao, Z., Wen, P.-C., Yoshioka, C., Tajkhorshid, E., and Gouaux, E. (2019) Serotonin transporter–ibogaine complexes illuminate mechanisms of inhibition and transport. Nature 569, 141.

37. Zhang, Y.-W., and Rudnick, G. (2006) The cytoplasmic substrate permeation pathway of serotonin transporter. Journal of Biological Chemistry 281, 36213–36220.

38. Merkle, P. S., Gotfryd, K., Cuendet, M. A., Leth-Espensen, K. Z., Gether, U., Loland, C. J., and Rand, K. D. (2018) Substrate-modulated unwinding of transmembrane helices in the NSS transporter LeuT. Science Advances 4, eaar6179.

39. Krishnamurthy, H., and Gouaux, E. (2012) X-ray structures of LeuT in substrate-free outward-open and apo inward-open states. Nature 481, 469–474.

40. Androutsellis-Theotokis, A., and Rudnick, G. (2002) Accessibility and conformational coupling in serotonin transporter predicted internal domains. Journal of Neuroscience 22, 8370–8378.

41. Zhang, Y.-W., and Rudnick, G. (2005) Cysteine-scanning mutagenesis of serotonin transporter intracellular loop 2 suggests an *α*-helical conformation. Journal of Biological Chemistry 280, 30807–30813.

42. Jacobs, M. T., Zhang, Y.-W., Campbell, S. D., and Rudnick, G. (2007) Ibogaine, a noncompetitive inhibitor of serotonin transport, acts by stabilizing the cytoplasm-facing state of the transporter. Journal of Biological Chemistry 282, 29441–29447.

43. Zhang, Y.-W., Tavoulari, S., Sinning, S., Aleksandrova, A. A., Forrest, L. R., and Rudnick, G. (2018) Structural elements required for coupling ion and substrate transport in the neurotransmitter transporter homolog LeuT. Proceedings of the National Academy of Sciences 115, E8854–E8862.

44. Zhao, Y., Terry, D. S., Shi, L., Quick, M., Weinstein, H., Blanchard, S. C., and Javitch, J. A. (2011) Substrate-modulated gating dynamics in a Na+-coupled neurotransmitter transporter homologue. Nature 474, 109–113.

45. Cheng, M. H., and Bahar, I. (2015) Molecular mechanism of dopamine transport by human dopamine transporter. Structure 23, 2171–2181.

46. Dehnes, Y., Shan, J., Beuming, T., Shi, L., Weinstein, H., and Javitch, J. A. (2014) Conformational changes in dopamine transporter intracellular regions upon cocaine binding and dopamine translocation. Neurochemistry international 73, 4–15.

47. Prasad, H. C., Zhu, C.-B., McCauley, J. L., Samuvel, D. J., Ramamoorthy, S., Shelton, R. C., Hewlett, W. A., Sutcliffe, J. S., and Blakely, R. D. (2005) Human serotonin transporter variants display altered sensitivity to protein kinase G and p38 mitogen-activated protein kinase. Proceedings of the National Academy of Sciences 102, 11545–11550.

48. Veenstra-VanderWeele, J. et al. (2012) Autism gene variant causes hyperserotonemia, serotonin receptor hypersensitivity, social impairment and repetitive behavior. Proceedings of the National Academy of Sciences 109, 5469–5474.

49. Quinlan, M. A., Krout, D., Katamish, R. M., Robson, M. J., Nettesheim, C., Gresch, P. J., Mash, D. C., Henry, L. K., and Blakely, R. D. (2019) Human Serotonin Transporter Coding Variation Establishes Conformational Bias with Functional Consequences. ACS Chemical Neuroscience 10, 3249–3260.

50. Annamalai, B., Mannangatti, P., Arapulisamy, O., Shippenberg, T. S., Jayanthi, L. D., and Ramamoorthy, S. (2011) Tyrosine Phosphorylation of the Human Serotonin Transporter: A Role in the Transporter Stability and Function. Molecular Pharmacology 81, 73–85.

51. Sørensen, L., Strømgaard, K., and Kristensen, A. S. (2014) Characterization of Intracellular Regions in the Human Serotonin Transporter for Phosphorylation Sites. ACS Chemical Biology 9, 935–944.

52. Penmatsa, A., Wang, K. H., and Gouaux, E. (2013) X-ray structure of dopamine transporter elucidates antidepressant mechanism. Nature 503, 85–90.

53. Zeppelin, T., Ladefoged, L. K., Sinning, S., Periole, X., and Schiøtt, B. (2018) A direct interaction of cholesterol with the dopamine transporter prevents its out-to-inward transition. PLoS computational biology 14, e1005907.

54. Laursen, L., Severinsen, K., Kristensen, K. B., Periole, X., Overby, M., Müller, H. K., Schiøtt, B., and Sinning, S. (2018) Cholesterol binding to a conserved site modulates the conformation, pharmacology, and transport kinetics of the human serotonin transporter. Journal of Biological Chemistry 293, 3510–3523.

55. Hamilton, P. J., Belovich, A. N., Khelashvili, G., Saunders, C., Erreger, K., Javitch, J. A., Sitte, H. H., Weinstein, H., Matthies, H. J., and Galli, A. (2014) PIP 2 regulates psychostimulant behaviors through its interaction with a membrane protein. Nature chemical biology 10, 582–589.

56. Khelashvili, G., Stanley, N., Sahai, M. A., Medina, J., LeVine, M. V., Shi, L., De Fabritiis, G., and Weinstein, H. (2015) Spontaneous inward opening of the dopamine transporter is triggered by PIP2-regulated dynamics of the N-terminus. ACS chemical neuroscience 6, 1825–1837.

57. Bartholomäus, I., Milan-Lobo, L., Nicke, A., Dutertre, S., Hastrup, H., Jha, A., Gether, U., Sitte, H. H., Betz, H., and Eulenburg, V. (2008) Glycine transporter dimers: evidence for occurrence in the plasma membrane. Journal of Biological Chemistry 283, 10978–10991.

58. Scholze, P., Freissmuth, M., and Sitte, H. H. (2002) Mutations within an intramembrane leucine heptad repeat disrupt oligomer formation of the rat GABA transporter 1. Journal of Biological Chemistry 277, 43682–43690.

59. Sitte, H. H., Farhan, H., and Javitch, J. A. (2004) Sodium-dependent neurotransmitter transporters: oligomerization as a determinant of transporter function and trafficking. Molecular interventions 4, 38.

60. Anderluh, A., Klotzsch, E., Reismann, A. W., Brameshuber, M., Kudlacek, O., Newman, A. H., Sitte, H. H., and Schütz, G. J. (2014) Single molecule analysis reveals coexistence of stable serotonin transporter monomers and oligomers in the live cell plasma membrane. Journal of Biological Chemistry 289, 4387–4394.

61. Cheng, M. H., Garcia-Olivares, J., Wasserman, S., DiPietro, J., and Bahar, I. (2017) Allosteric modulation of human dopamine transporter activity under conditions promoting its dimerization. Journal of Biological Chemistry 292, 12471–12482.

62. Periole, X., Zeppelin, T., and Schiøtt, B. (2018) Dimer interface of the human serotonin transporter and effect of the membrane composition. Scientific reports 8, 1–15.

63. Jayaraman, K., Morley, A. N., Szöllősi, D., Wassenaar, T. A., Sitte, H. H., and Stockner, T. (2018) Dopamine transporter oligomerization involves the scaffold domain, but spares the bundle domain. PLoS computational biology 14, e1006229.

64. Möller, I. R., Slivacka, M., Nielsen, A. K., Rasmussen, S. G., Gether, U., Loland, C. J., and Rand, K. D. (2019) Conformational dynamics of the human serotonin transporter during substrate and drug binding. Nature communications 10, 1–13.

65. Billesbølle, C. B., Mortensen, J. S., Sohail, A., Schmidt, S. G., Shi, L., Sitte, H. H., Gether, U., and Loland, C. J. (2016) Transition metal ion FRET uncovers K+ regulation of a neurotransmitter/sodium symporter. Nature communications 7, 1–12.

66. Coleman, J. A., and Gouaux, E. (2018) Structural basis for recognition of diverse antidepressants by the human serotonin transporter. Nature structural & molecular biology 25, 170–175.

67. Trott, O., and Olson, A. J. (2010) AutoDock Vina: improving the speed and accuracy of docking with a new scoring function, efficient optimization, and multithreading. Journal of computational chemistry 31, 455–461.

68. Wang, K. H., Penmatsa, A., and Gouaux, E. (2015) Neurotransmitter and psychostimulant recognition by the dopamine transporter. Nature 521, 322–327.

69. Jorgensen, W. L., Chandrasekhar, J., Madura, J. D., Impey, R. W., and Klein, M. L. (1983) Comparison of simple potential functions for simulating liquid water. The Journal of chemical physics 79, 926–935.

70. Maier, J. A., Martinez, C., Kasavajhala, K., Wickstrom, L., Hauser, K. E., and Simmerling, C. (2015) ff14SB: improving the accuracy of protein side chain and backbone parameters from ff99SB. Journal of chemical theory and computation 11, 3696–3713.

71. Wang, J., Wang, W., Kollman, P. A., and Case, D. A. (2006) Automatic atom type and bond type perception in molecular mechanical calculations. Journal of molecular graphics and modelling 25, 247–260.

72. Case, D. A. et al. (2018) AMBER 2018. University of California, San Francisco

73. Jo, S., Kim, T., Iyer, V. G., and Im, W. (2008) CHARMM-GUI: a web-based graphical user interface for CHARMM. Journal of computational chemistry 29, 1859–1865.

74. Huang, J., Rauscher, S., Nawrocki, G., Ran, T., Feig, M., de Groot, B. L., Grubmüller, H., and MacKerell, A. D. (2017) CHARMM36m: an improved force field for folded and intrinsically disordered proteins. Nature methods 14, 71–73.

75. Krutler, V., Gunsteren, W. F. v., and Hnenberger, P. H. (2001) A fast SHAKE algorithm to solve distance constraint equations for small molecules in molecular dynamics simulations. Journal of Computational Chemistry 22, 501–508.

76. York, D. M., Darden, T. A., and Pedersen, L. G. (1993) The effect of longrange electrostatic interactions in simulations of macromolecular crystals: A comparison of the Ewald and truncated list methods. The Journal of Chemical Physics 99, 8345–8348.

77. Lee, T.-S., Cerutti, D. S., Mermelstein, D., Lin, C., LeGrand, S., Giese, T. J., Roitberg, A., Case, D. A., Walker, R. C., and York, D. M. (2018) GPU-accelerated molecular dynamics and free energy methods in Amber18: performance enhancements and new features. Journal of chemical information and modeling 58, 2043–2050.

78. Eastman, P., Swails, J., Chodera, J. D., McGibbon, R. T., Zhao, Y., Beauchamp, K. A., Wang, L.-P., Simmonett, A. C., Harrigan, M. P., Stern, C. D., Wiewiora, R. P., Brooks, B. R., and Pande, V. S. (2017) OpenMM 7: Rapid development of high performance algorithms for molecular dynamics. PLOS Computational Biology 13, e1005659.

79. Wang, J., Wolf, R. M., Caldwell, J. W., Kollman, P. A., and Case, D. A. (2004) Development and testing of a general amber force field. Journal of computational chemistry 25, 1157–1174.

80. Hopkins, C. W., Le Grand, S., Walker, R. C., and Roitberg, A. E. (2015) Long-time-step molecular dynamics through hydrogen mass repartitioning. Journal of chemical theory and computation 11, 1864–1874.

81. Husic, B. E., and Pande, V. S. (2018) Markov state models: From an art to a science. Journal of the American Chemical Society 140, 2386–2396.

82. Shukla, D., Hernández, C. X., Weber, J. K., and Pande, V. S. (2015) Markov State Models Provide Insights into Dynamic Modulation of Protein Function. 48, 414–422.

83. Chodera, J. D., and Noé, F. (2014) Markov state models of biomolecular conformational dynamics. 25, 135–144.

84. Gu, S., Silva, D.-A., Meng, L., Yue, A., and Huang, X. (2014) Quantitatively Characterizing the Ligand Binding Mechanisms of Choline Binding Protein Using Markov State Model Analysis. 10, e1003767.

85. Zimmerman, M. I. et al. (2021) SARS-CoV-2 simulations go exascale to predict dramatic spike opening and cryptic pockets across the proteome. Nature Chemistry 13, 651–659.

86. Moffett, A. S., and Shukla, D. (2018) Using molecular simulation to explore the nanoscale dynamics of the plant kinome. Biochemical Journal 475, 905–921.

87. Selvam, B., Mittal, S., and Shukla, D. (2018) Free Energy Landscape of the Complete Transport Cycle in a Key Bacterial Transporter. ACS Central Science 4, 1146–1154.

88. Selvam, B., Yu, Y.-C., Chen, L.-Q., and Shukla, D. (2019) Molecular Basis of the Glucose Transport Mechanism in Plants. ACS Central Science 5, 1085–1096.

89. Cheng, K. J., Selvam, B., Chen, L.-Q., and Shukla, D. (2019) Distinct substrate transport mechanism identified in homologous sugar transporters. The Journal of Physical Chemistry B 123, 8411–8418.

90. Feng, J., Selvam, B., and Shukla, D. (2021) How do antiporters exchange substrates across the cell membrane? An atomic-level description of the complete exchange cycle in NarK. 29, 922–933.e3.

91. Chan, M. C., and Shukla, D. (2021) Markov state modeling of membrane transport proteins. Journal of Structural Biology 213, 107800.

92. Meng, Y., Shukla, D., Pande, V. S., and Roux, B. (2016) Transition path theory analysis of c-Src kinase activation. 113, 9193–9198.

93. Roe, D. R., and Cheatham, T. E. (2013) PTRAJ and CPPTRAJ: Software for Processing and Analysis of Molecular Dynamics Trajectory Data. Journal of Chemical Theory and Computation 9, 3084–3095.

94. McGibbon, R., Beauchamp, K., Harrigan, M., Klein, C., Swails, J., Hernndez, C., Schwantes, C., Wang, L.-P., Lane, T., and Pande, V. (2015) MDTraj: A Modern Open Library for the Analysis of Molecular Dynamics Trajectories. Biophysical Journal 109, 1528–1532.

95. Scherer, M. K., Trendelkamp-Schroer, B., Paul, F., Prez-Hernndez, G., Hoffmann, M., Plattner, N., Wehmeyer, C., Prinz, J.-H., and No, F. (2015) PyEMMA 2: A Software Package for Estimation, Validation, and Analysis of Markov Models. Journal of Chemical Theory and Computation 11, 5525–5542.

96. Humphrey, W., Dalke, A., and Schulten, K. (1996) VMD: Visual molecular dynamics. Journal of Molecular Graphics 14, 33–38.

