## Supplementary Methods, Images, Tables and Results for "Structural rearrangement of the intracellular gate of the serotonin transporter induced by Thr276 phosphorylation"

### Supporting information for Structural rearrangement of the intracellular gate of the serotonin transporter induced by Thr276 phosphorylation

Matthew C. Chan,[†] Erik Procko,[‡,¶,§,‖] and Diwakar Shukla[*,†,¶,‖,⊥,#,@]

†*Department of Chemical and Biomolecular Engineering, University of Illinois at Urbana-Champaign, Urbana, IL, 61801*

‡*Department of Biochemistry, University of Illinois at Urbana-Champaign, Urbana, IL, 61801*

¶*Center for Biophysics and Quantitative Biology, University of Illinois at Urbana-Champaign, Urbana, IL, 61801*

§*Neuroscience Program, University of Illinois at Urbana-Champaign, Urbana, IL, 61801*

‖*Cancer Center at Illinois, University of Illinois at Urbana-Champaign, Urbana, IL, 61801*

⊥*National Center for Supercomputing Applications, University of Illinois, Urbana, IL, 61801*

#*Beckman Institute for Advanced Science and Technology, University of Illinois at Urbana-Champaign, Urbana, IL, 61801*

@*NIH Center for Macromolecular Modeling and Bioinformatics, University of Illinois at Urbana-Champaign, Urbana, IL, 61801*

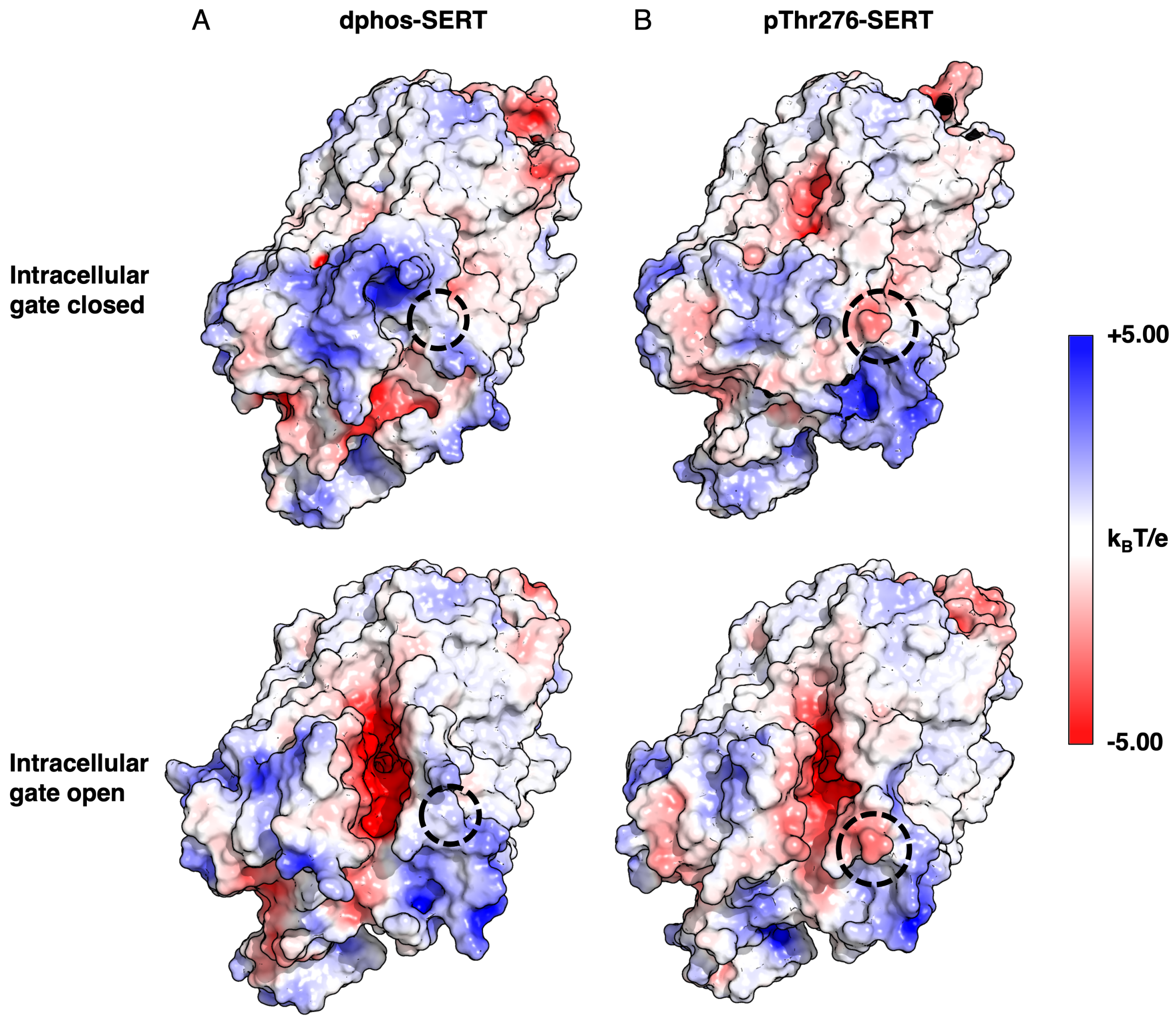


Figure S1: Electrostatic potential map of WT-SERT (A) and pThr276-SERT (B) viewed from the intracellular side. Electrostatic potential is colored from negatively charged (red) to neutral (white) to positively charged (blue). The Thr276 phosphorylation site is circled.

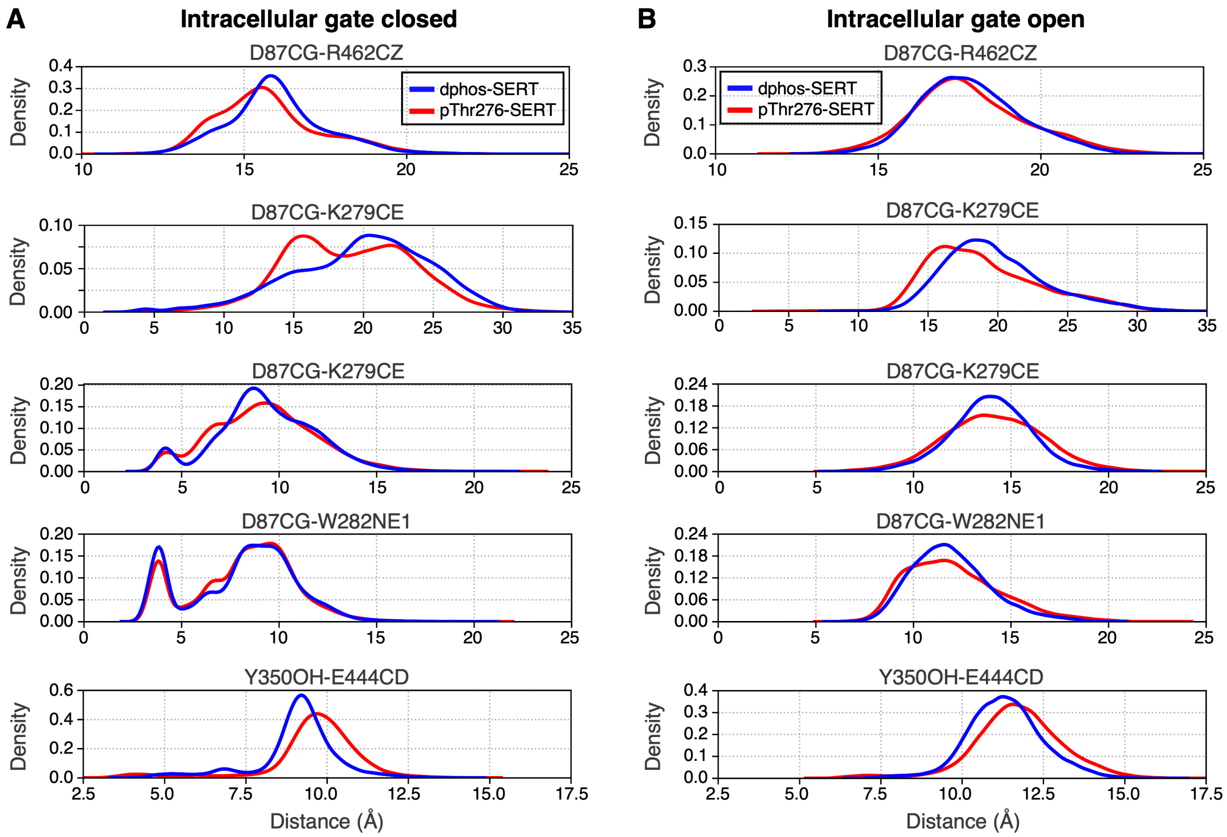


Figure S2: Distance distribution of select gating residues for 50,000 MD structures for each occluded (A) and inward-facing (B). Distances calculated from the WT-SERT MD simulations is represented in blue while pThr276-SERT in red.

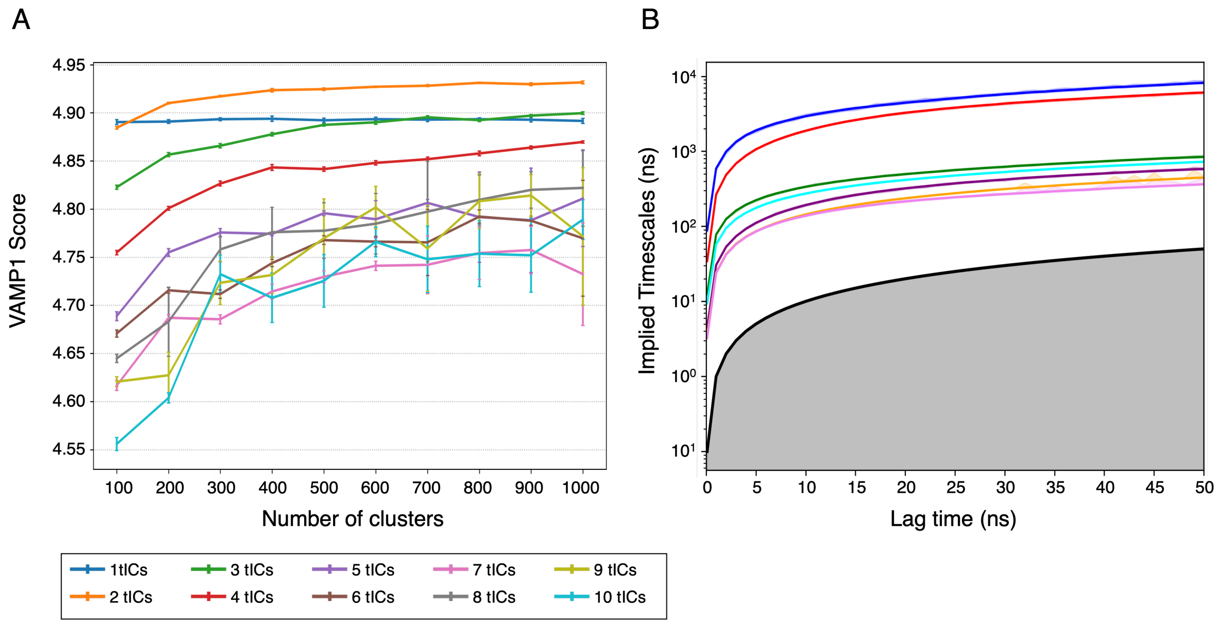


Figure S3: (A) Optimization of number of MSM clusters and time-independent components (tICs) based on maximizing the VAMP1 score. The phosphorylated SERT MSM was constructed using 500 clusters and 2 tICs. (B) Implied timescale plots from the transition probability matrix of the MSM. The first seven eigenvalues corresponding to the seven slowest transition rates obtained from the constructured MSM using 500 clusters and 2 tICs is plotted. The Markovian lag time was chosen as 12 ns.

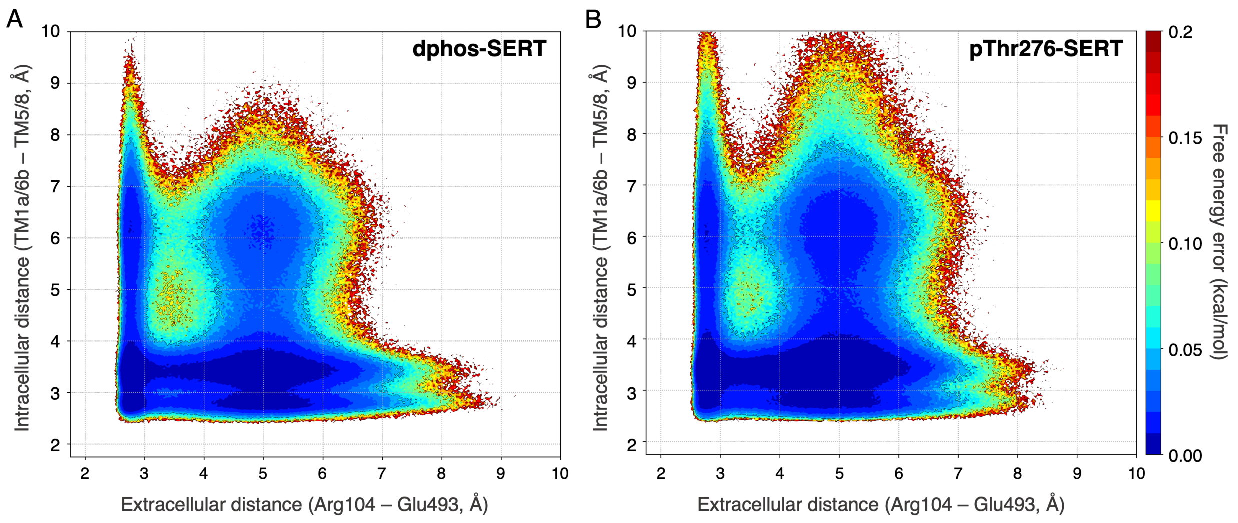


Figure S4: Standard error of the free energy landscape projected on the coordinates of the extracellular and intracellular gating distances of SERT. The error was calculated with the bootstrapping method by constructing a MSM with randomly selecting 80% of the trajectory data for 500 independent samples.

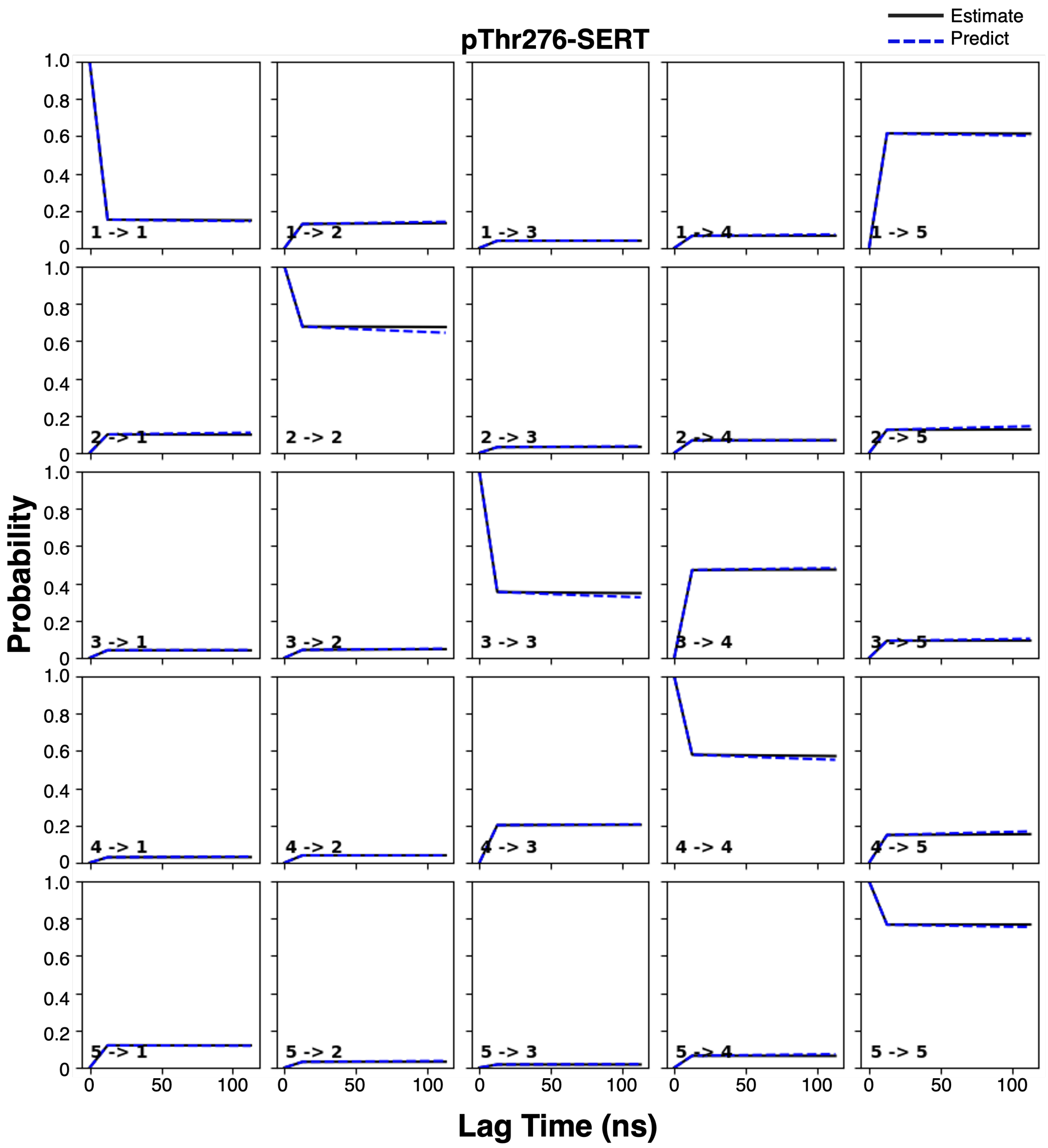


Figure S5: Chapman-Kolmogorov validation of the pThr276-SERT Markov state model. The Chapman-Kolmogorov test was implemented using the pyEMMA package and performed on 5 macrostates of the Markov model.
